# The Theory and Practice of the viral dose in neutralization assay: insights on SARS-CoV-2 “doublethink” effect

**DOI:** 10.1101/2020.10.16.342428

**Authors:** Alessandro Manenti, Eleonora Molesti, Marta Maggetti, Alessandro Torelli, Giulia Lapini, Emanuele Montomoli

## Abstract

Due to the global spread of the severe acute respiratory syndrome coronavirus 2 (SARS-CoV-2), there is an urgent need for reliable high-throughput serological assays in order to evaluate the immunological responses against SARS-COV-2 virus and to enable population screening, as well as vaccines and drug’s efficacy testing. Several serological assays for SARS-CoV-2 are now becoming available in the market. However, it has also become extremely important to have well-established assays with desirable high sensitivity and specificity. To date, the micro-neutralization (MN) assay, is currently considered the gold-standard being capable of evaluating and detecting, functional neutralizing antibodies (nAbs). Several protocols exist for microneutralization assays which vary in several steps of the protocol: cell seeding conditions, number of cells seeded, virus amount used in the infection step, virus-serum-cells incubation period etc. These potential differences account for a high degree of variability and inconsistency of the results and using a harmonized protocol for the micro-neutralization assay could potentially solve this.

Given this situation, the main aim of our study was to carry out SARS-CoV-2 wild type virus MN assay in order to investigate which optimal tissue culture infective dose 50 (TCID_50_) infective dose in use is the most adequate choice for implementation in terms of reproducibility, standardization possibilities and comparability of results. Therefore, we assessed the MN by using two different viral infective doses: a standard dose of 100 TCID_50_/well and a lower dose of 25 TCID_50_/well. The results obtained, yielded by MN on using the lower infective dose (25 TCID_50_), were in line with those obtained with the standard infective dose; in some cases, however, we detected a titre that was one or two dilution steps higher, which maintained all negative samples negative. This suggesting that the lower dose can potentially have a positive impact on the detection and estimation of neutralizing antibodies present in a given sample, showing higher sensitivity but similar specificity and therefore, it would require a more accurate assessment and cross-laboratories standardisation especially when MN is employed as serological assay of choice for pre-clinical and clinical studies.

## Introduction

The detection and quantitation of serum antibodies to different viral antigens, after natural infection and/or immunization, has long been used to assess the likelihood of protection against a specific pathogen [1]. The Enzyme Linked-immunosorbent assay (ELISA) is one of the most used method for total antibodies detection. This method is able to detect all the antibodies present in a given sample able to bind the specific antigen of interest coated in a dedicated plate. It is fast, cheap and safe because it does not require the handling of live pathogens. Another classical way of measuring antibodies for agglutinating viruses such as Influenza virus, is the Haemagglutination inhibition assay (HAI). This method, considered as the gold standard in Influenza field [2,3]. It is based upon the principle that antibody binding to the HA globular head can inhibit the HA’s ability to agglutinate red blood cells (RBCs) by preventing the binding between the head domain (HA1) of the hemagglutinin (HA) and the sialic acids (SA) present on the RBC surface. Both, ELISA and HAI suffer from the fact that they are not able to give a precise indication about the functionality of the antibodies detected. Given these limitations, the Micro-Neutralization assay (MN) is an attractive alternative for the assessment of baseline serostatus and the evaluation of the humoral responses following natural infection and/or vaccination [4]. MN assays were developed in 1990 [5,6]. This is a functional assay, and it is able to detect neutralizing antibodies capable of prevent the virus infection of different mammalian cell lines and the neutralization activity is measured as the ability of the sera to reduce the CPE due to inhibition of viral entry and subsequent replication [7]. Compared to the ELISA-based methods, the results derived by the MN represent a more precise and relevant estimation of antibody-mediated protection in-vitro [8].

On the other hand, MN is more complex to perform due to some requirements of the test itself: the need of live viruses and biosecurity level 4, 3 or 2+ laboratories in case of class IV, III or II pathogens; it is more expensive; and there are difficulties in protocol standardization across laboratories (e.g. cell lines, infective dose, days of incubation and read-out).

In the present study we focused our attention on the performance of the MN assay with SARS-CoV-2 wild type virus using two different input of viral dose: the standard 100 Tissue Culture Infective Dose 50% (TCID_50_) and the 25 TCID_50_ infective dose. The data obtained were than used to investigate the optimal viral dose that should be effectively used for SARS-CoV-2 strain in the MN assay.

## Methods

### Serum samples and human monoclonal antibody IgG1

A total of 102 human serum samples were collected as part of a sero-epidemiological study that is being performed in the laboratory of Molecular Epidemiology of the University of Siena, Italy [9]. Serum samples were anonymously collected in compliance with Italian ethics law. The human monoclonal antibody IgG1-CR3022 (Absolute Antibody) and the human monoclonal antibody IgG1 SAD-S35 (Acrobiosystem) were tested along with the serum samples in the MN assay and ELISA. Hyperimmune sheep antisera against Influenza A/H1N1/California/7/2009 (10/218), B/Brisbane/60/2008 (13/312) and A/Anhui/1/2013 (15/248) strains were purchased from the National Institute for Biological Standard and Controls (NIBSC, UK). Hyperimmune rabbit serum samples against Adenovius Type 4 (V204-502-565) were provided by National Institute of Allergy and Infectious Diseases (NIH, Bethesda). Human serum minus IgA/IgM/IgG (S5393-1VL) (Sigma, St. Louis, MO, USA) was used as a negative control.

### Cell cultures

VEROE6 cells, an epithelial cell line from the kidney of a normal monkey (Cercopithecus aethiops), were acquired from the American Type Culture Collection (ATCC – CRL 1586).

VEROE6 cells were cultured in Dulbecco’s Modified Eagle’s Medium (DMEM) – High Glucose (Euroclone, Pero, Italy) supplemented with 2 mM L-Glutamine (Lonza, Milano, Italy), 100 units/mL penicillinstreptomycin mixture (Lonza, Milano, Italy) and 10% of foetal bovine serum (FBS) (Euroclone, Pero, Italy) at 37°C, in a 5% CO2 humidified incubator.

Adherent sub-confluent cell monolayers of VERO E6 were prepared in growth medium, D-MEM high glucose containing 2% FBS in 96-well plates for titration and neutralization tests of SARS-CoV-2.

### Virus and Titration

SARS CoV-2 2019-nCov/Italy-INMI1 – wild type virus was purchased from the European Virus Archive goes Global (EVAg, Spallanzani Institute, Rome). The virus was titrated in serial 1 log dilutions (from 1 log to 11 log) to obtain a 50% tissue culture infective dose (TCID_50_) on 96-well culture plates of VERO and VERO E6 cells. The plates were observed daily for a total of 4 days for the presence of CPE by means of an inverted optical microscope. The end-point titres were calculated according to the Reed & Muench method based on eight replicates for titration as described before [10].

### Micro-neutralization assay and CPE-Read Out

Serum samples were heat-inactivated for 30 minutes at 56°C; 2-fold serial dilutions, starting from 1:10, were then mixed with an equal volume of viral solution containing 100 and 25 TCID_50_ of SARS-CoV-2. The serum-virus mixture was incubated for 1 hour at 37°C in a humidified atmosphere with 5% CO2. After incubation, 100 μl of the mixture at each dilution was added in duplicate to a cell plate containing a semi-confluent VERO E6 monolayer. The plates were incubated for 72±8 hours (H) at 37°C in a humidified atmosphere with 5% CO2.

After three days of incubation, the plates were inspected by an inverted optical microscope. The highest serum dilution that protected more than the 50% of cells from CPE was taken as the neutralization titre.

### Enzyme-Linked Immunosorbent Assay (ELISA)

Specific anti-SARS-CoV-2 IgG antibodies were detected through a commercial ELISA kit (Euroimmun, Lübeck, Germany). ELISA plates are coated with recombinant structural protein (S1 domain) of SARS-CoV-2. According to the manufacturer, cross-reactions may occur with anti-SARS-CoV(−1) IgG antibodies, due to the close relationship between SARS-CoV(−1) and SARS-CoV-2, while cross-reactions with other human pathogenic CoVs (MERS-CoV, HCoV-229E, HCoV-NL63, HCoV-HKU1, and HCoV-OC43) are excluded. The assay provides semi-quantitative results by calculating the ratio of the OD of the serum sample over the OD of the calibrator. According to the manufacturer’s instructions, positive samples have a ratio ≥ 1.1, borderline samples a ratio between 0.8 and 1.1 and negative samples a ratio < 0.8.

### Statistics analysis

Data analysis was performed using GraphPad Prism Version 5. The Kruskal-Wallis non-parametric test [11] was used to evaluate the difference between MN performed with 100 and 25 TCID_50_ titres.

## DISCUSSIONS AND CONCLUSIONS

Among different serological tests, the MN is the only assay that can offer a high throughput in processing samples along with the information regarding the ability of the antibodies to prevent the virus attachment and enter into the target cells. To date, MN assay is considered the reference standard method for detection of neutralizing antibodies, which may be used as a correlate of protective immunity. Although alternative BSL2 protocols using SARS CoV-2 pseudotyped viruses are being developed to obviate culture of live SARS-CoV-2 [12–14] these methods remain in the research area.

Historically, such as for Influenza virus, the MN assay is routinely carried out in 96-micro-well plates, by mixing different 2-fold serial dilutions of a serum-containing antibodies with a well-defined viral dose containing 100 TCID_50_/well. However, for newly emerging viruses such as SARS-CoV-2, the viral dose needs to be accurately evaluated necessitating agreement on a consensus assay protocol for future studies.

The viral load equal to 100 TCID_50_, in accordance with the empirical formula obtained by applying the Poisson distribution, should be equal to approximately 70 plaque-forming units (pfu), which represents the measure of the infectious viral particles in a certain volume of medium used in each well of the microplate. Clearly, this is valid if the same cell system is used and the virus is able to form plaques on the cells monolayer.

In this preliminary study, we evaluated the results obtained by MN CPE-based test with wild type SARS-CoV-2 pandemic strain, using two different viral load in the assay: 100 and 25 TCID_50_/well.

All the 102 serum samples screened have been assayed by ELISA test in order to assess more specifically the presence/absence of anti-SARS-CoV-2 binding antibodies. Among the ELISA positive sample 19.8-20 % of sera were found positive in MN assay with 100 TCID50 and 25 TCID50 of viral dose.

According to the results obtained and, based on the assumption that there are not defined indications of the viral dose required for functional assays such as MN or plaque reduction, along with the fact that 100 TCID_50_ is the viral load used for other respiratory viruses such as Influenza, we tested the same sample panel by using an even lower infection dose: 25 TCID50.

Our results show that, with the lower dose (25 TCID_50_), in the majority of the cases the MN titres are higher of one or two dilution steps (Table 1, Figure 1). More interestingly, all the negative MN 100TCID_50_ serum samples were also confirmed negative by MN 25TCID_50_.

**Table 1.** ELISA and Neutralization results for all 102 human serum samples. Negative samples are indicated in the first row of the table. Neutralizing titres, obtained with CPE (100 and 25 TCID_50_ infective dose) are indicated for each sample

| *Sample ID* | **ELISA** | **MN CPE Titre**<br>**100 TCID<sub>50</sub>** | **MN CPE Titre**<br>**25 TCID<sub>50</sub>** |
| --- | --- | --- | --- |
| *From 1 to 28* | Negative | 5 | 5 |
| *From 29 to 45* | Borderline | 5 | 5 |
| **46** | Borderline | 20 | 40 |
| **47** | Borderline | 80 | 80 |
| **48** | Borderline | 20 | 20 |
| **49** | Positive | 640 | 640 |
| **50** | Positive | 20 | 40 |
| **51** | Positive | 320 | 320 |
| **52** | Positive | 640 | 640 |
| **53** | Positive | 40 | 40 |
| **54** | Positive | 640 | 640 |
| **55** | Positive | 20 | 20 |
| **56** | Positive | 10 | 20 |
| **57** | Positive | 160 | 320 |
| **58** | Positive | 40 | 40 |
| **59** | Positive | 160 | 320 |
| **60** | Positive | 640 | 640 |
| **61** | Positive | 80 | 80 |
| **62** | Positive | 1280 | 1280 |
| **63** | Positive | 160 | 320 |
| **64** | Positive | 80 | 80 |
| **65** | Positive | 10 | 20 |
| **66** | Positive | 80 | 80 |
| **67** | Positive | 640 | 640 |
| **68** | Positive | 10 | 10 |
| **69** | Positive | 40 | 40 |
| **70** | Positive | 40 | 40 |
| **71** | Positive | 40 | 160 |
| **72** | Positive | 40 | 80 |
| **73** | Positive | 20 | 80 |
| **74** | Positive | 40 | 160 |
| **75** | Positive | 10 | 20 |
| **76** | Positive | 40 | 80 |
| **77** | Positive | 5 | 20 |
| **78** | Positive | 20 | 40 |
| **79** | Positive | 40 | 160 |
| **80** | Positive | 10 | 40 |
| **81** | Positive | 40 | 160 |
| **82** | Positive | 320 | 640 |
| *From 83 to 102* | Positive | 5 | 5 |

| *Sample ID* | **ELISA** | **MN CPE Titre**<br>**100 TCID<sub>50</sub>** | **MN CPE Titre**<br>**25 TCID<sub>50</sub>** |
| --- | --- | --- | --- |
| ***From 1 to 28*** | Negative | 5 | 5 |
| ***From 29 to 45*** | Borderline | 5 | 5 |
| **46** | Borderline | 20 | 40 |
| **47** | Borderline | 80 | 80 |
| **48** | Borderline | 20 | 20 |
| **49** | Positive | 640 | 640 |
| **50** | Positive | 20 | 40 |
| **51** | Positive | 320 | 320 |
| **52** | Positive | 640 | 640 |
| **53** | Positive | 40 | 40 |
| **54** | Positive | 640 | 640 |
| **55** | Positive | 20 | 20 |
| **56** | Positive | 10 | 20 |
| **57** | Positive | 160 | 320 |
| **58** | Positive | 40 | 40 |
| **59** | Positive | 160 | 320 |
| **60** | Positive | 640 | 640 |
| **61** | Positive | 80 | 80 |
| **62** | Positive | 1280 | 1280 |
| **63** | Positive | 160 | 320 |
| **64** | Positive | 80 | 80 |
| **65** | Positive | 10 | 20 |
| **66** | Positive | 80 | 80 |
| **67** | Positive | 640 | 640 |
| **68** | Positive | 10 | 10 |
| **69** | Positive | 40 | 40 |
| **70** | Positive | 40 | 40 |
| **71** | Positive | 40 | 160 |
| **72** | Positive | 40 | 80 |
| **73** | Positive | 20 | 80 |
| **74** | Positive | 40 | 160 |
| **75** | Positive | 10 | 20 |
| **76** | Positive | 40 | 80 |
| **77** | Positive | 5 | 20 |
| **78** | Positive | 20 | 40 |
| **79** | Positive | 40 | 160 |
| **80** | Positive | 10 | 40 |
| **81** | Positive | 40 | 160 |
| **82** | Positive | 320 | 640 |
| ***From 83 to 102*** | Positive | 5 | 5 |

**Figure 1.**
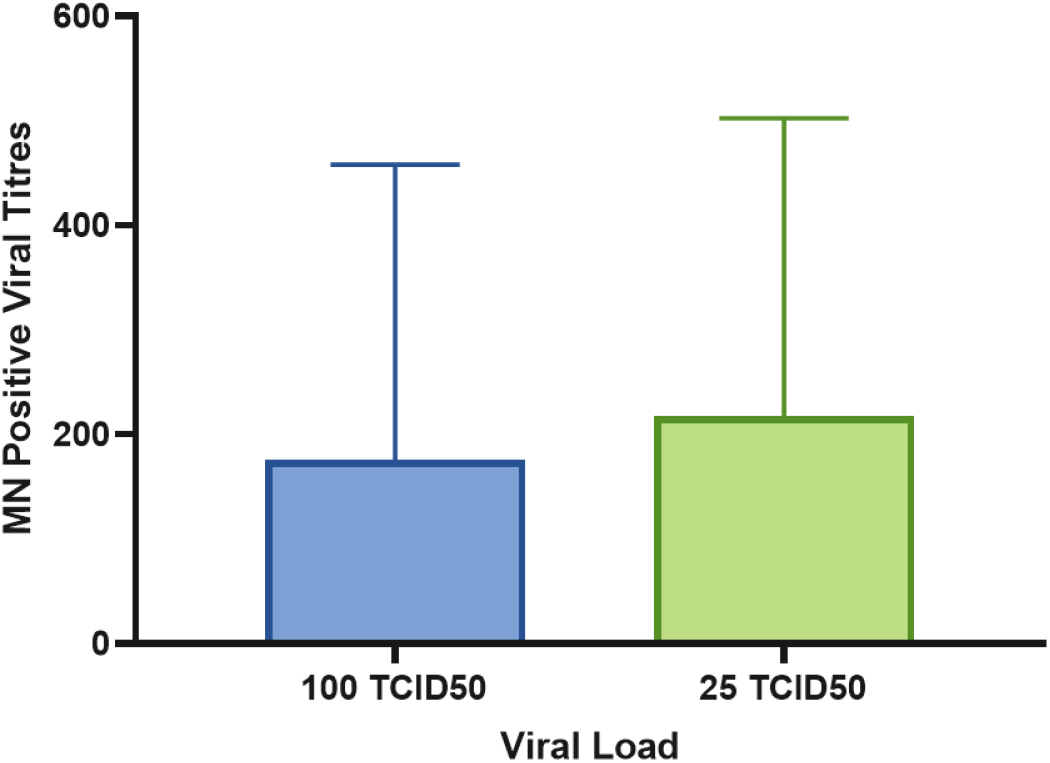
MN positive CPE-viral titres obtained when 102 samples were tested against 100 TCID_50_ and 25 TCID_50_ SARS-CoV-2 analysed by GraphPad using Kruskal-Wallis non-parametric test.

These results are of considerable importance leading to the conclusion that even if a lower infective dose is used, the possibility to have false positives in ELISA and MN 100 TCID_50_ confirmed-negative samples is low, and that the sensitivity of the assay to detect functional antibodies could be improved by reducing the viral dose. As also evidenced in figure 2, if no neutralizing antibodies are present no significant differences in the read-out (percentage of CPE present in the first dilution) are observed for both viral loads.

**Figure 2.**
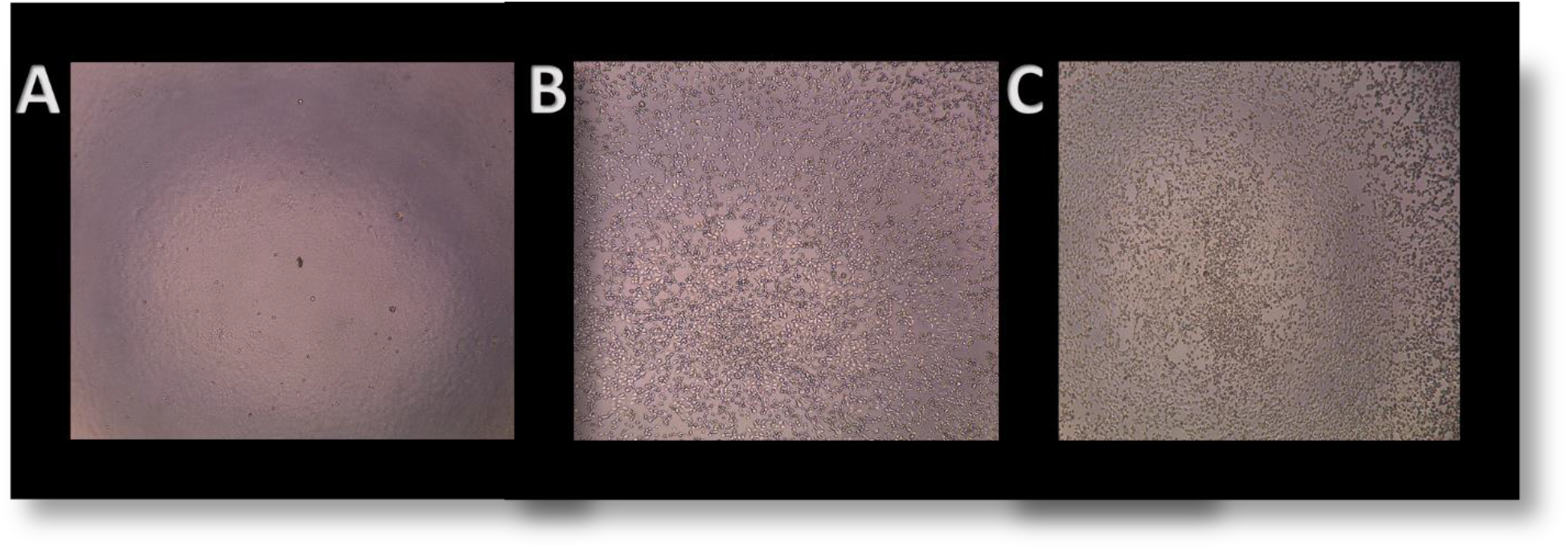
Vero E6 cells at different stage of infection. A) Not infected VERO E6 cell monolayer (cell control) complete absence of CPE; B) 1:10 dilution of a negative serum sample after 72 hours of incubation with 100 TCID_50_ infective dose of SARS-CoV-2; C) 1:10 dilution of a negative serum sample after 72 hours of incubation with 25 TCID_50_ infective dose of SARS-CoV-2.

Thus confirming that even with a lower infective dose the cell monolayer is able to results in 100 % CPE after 3 days (72h) of incubation, avoiding the possibility to have false positive results due to non-specific inhibition of the viral infection by the high serum concentration at the first sample dilution.

This aspect could be crucial in order to evaluate, in a specific manner, the immune response against new emerging viruses, such as the SARS-CoV-2, for which immunological and serological data need to be well established and defined. In fact, there is no doubt that the absence of oversight and standardisation of serologic tests is concerning given that the available serologic assays are highly variable, differing in their format, the antibody class detected, the selected antigen, and the acceptable sample types [15].

Notably, regardless of which neutralizing antibody test is being performed, it is still unclear what minimal neutralizing antibody titer correlates with protective immunity and whether results from the available SARS-CoV-2 serologic assays can predict such immunity.

As evidenced before [1,16], it is important to note that serological assays able to detect a neutralizing antibody response will be critical to provide the most accurate results for vaccine immunogenicity trials. Understanding of the proper viral dose to be used would definitely be of great importance for the successful development and licensing of new vaccines and therapeutics against Covid-19 disease [17].

As first step towards this, the use of the standard 100 TCID_50_ viral dose could be reduced in order to increase the sensitivity of the test for SARS-CoV-2 since no precise indications or protocols are established. Even if small and preliminary, this study aims to encourage further international collaborations towards the standardization of the SARS-CoV-2 MN assay.

